# Attention-based approach to predict drug-target interactions across seven target superfamilies

**DOI:** 10.1101/2024.03.20.585893

**Authors:** Aron Schulman, Juho Rousu, Tero Aittokallio, Ziaurrehman Tanoli

## Abstract

**Motivation:** Drug-target interactions (DTIs) hold pivotal role in drug repurposing and elucidation of drug mechanisms of action. While single-targeted drugs have demonstrated clinical success, they often exhibit limited efficacy against complex diseases, such as cancers, whose development and treatment is dependent on several biological processes. Therefore, a comprehensive understanding of primary, secondary, and even inactive targets becomes essential in the quest for effective and safe treatments for cancer and other indications. The human proteome offers over a thousand druggable targets. Yet, most FDA-approved drugs bind with only a small fraction of disease targets.

**Results:** This study introduces an attention-based method to predict drug-target bioactivities for all human proteins across seven superfamilies. Nine different descriptor sets were meticulously examined to identify optimal signature descriptors for predicting novel DTIs. Our testing results demonstrated Spearman correlations exceeding 0.72 (P<0.001) for six out of seven superfamilies. The proposed method outperformed nine state-of-the-art deep learning and graph-based methods, and importantly, maintained relatively high performance for most target superfamilies when tested in independent sources of bioactivity data. We computationally validated 185,676 drug-target pairs from ChEMBL-V33, that were not available in model training, with a reasonable performance of Spearman correlation > 0.57 (P<0.001) for most superfamilies. This justifies the robustness of the proposed method for predicting novel DTIs. Finally, we applied our method to predict missing activities among 3,492 approved molecules in ChEMBL-V33, offering a valuable tool for advancing drug mechanism discovery and repurposing existing drugs for new indications.

**Availability and implementation:** The codes and training datasets to reproduce and extend the results are available at https://github.com/AronSchulman/MMAtt-DTA.

## 1. INTRODUCTION

Developing novel pharmaceutical agents is a formidable endeavor, incurring expenditures amounting to billions of dollars and necessitating a protracted timeframe of 9–15 years for successful market introduction [1]. Consequently, the pharmaceutical industry is increasingly motivated to explore alternative avenues, such as identifying new therapeutic applications for existing approved drugs. This strategic approach, commonly termed drug repurposing, holds considerable appeal due to its potential to accelerate drug development, curtail associated costs, and address unmet medical needs [2]. Drug-target interactions (DTI), i.e the target molecules each compound binds to at a binding strength that impacts on cellular functions, lie at the heart of drug discovery and repurposing. ChEMBL [3], BindingDB [4], DrugBank [5], GtopDB [6], DrugTargetCommons [7] and DgiDB [8] are the six most comprehensive, manually curated drug-target databases. These databases meticulously curate experimental bioactivity data spanning millions of compounds and encompassing thousands of protein targets. Despite the wealth of information these resources offer, it is pertinent to observe that none of these repositories affords comprehensive coverage of target interactions for approved drugs at the holistic proteome level. Therefore, several deep learning-based methods are proposed as an alternative to fill this gap [9].

Attention-based methods, as extension of deep learning, have gained major interest in drug discovery due to the breakthrough of the Transformers [10] in the field of natural language processing. In DTI predictions, the use of attention ranges from learning numerical descriptors for compounds and targets to contextualizing their complex relationships. MolTrans [11] is among the first methods to utilize parts of the Transformer architecture in DTI prediction. Specifically, MolTrans generates descriptors by applying an attention-based mechanism on one-hot encoded compound and target substructures. MolTrans only operates on the binary classification regime. While MolTrans remains a strong benchmarking model, several approaches have improved its limitations and design choices. Kang et al. [12] introduced a regression method for predicting continuous binding affinity values. Their method generates descriptors via transfer learning from pre-trained Bidirectional Encoder Representation from Transformers (BERT) based methods [13], where compound’s SMILES string and protein’s amino acid (AA) sequences are treated analogously to letters and words in natural language. The use of pre-trained methods allows access to vast amounts of unlabelled data. Subsequently, TransDTI [14] explores a wider variety of pretrained methods for proteins and establishes embeddings generated by ESM [15] and AlphaFold [16] as good alternatives for BERT.

Some of the existing methods do not restrict their use of attention to only generating embeddings. DTITR [17] and MCANet [18] successfully demonstrated how cross-attention can be applied across the descriptors of protein targets and compounds for improved interaction predictions. Nevertheless, these methods rely solely on SMILES and AA sequences to generate descriptors for compounds and protein targets, respectively. The availability and ease of use of such data is a practical advantage, but they can be viewed as overly simplified descriptors unable to capture all intricacies of structure and function. Some methods, such as Multi-TransDTI [19], utilize additional descriptors alongside Transformer-generated descriptors. Yet, they still perform the prediction task with simplistic concatenations and fully connected layers without applying attention. As a further alternative, MHTAN-DTI [20] and DTI-GTN [21] include interactions between drugs, proteins, diseases and side effects in a graph format. They utilize attention mechanisms tailored to suit graphs for promising DTI prediction results.

This work presents attention-based methods for DTI predictions in a regression setting for each of the seven superfamilies of the protein targets: enzymes, kinases, GPCRs, nuclear receptors, epigenetic receptors, ion channels and transporters. Similar to the current state-of-the-art, our methods utilize attention for generating compound and protein descriptors. However, instead of directly encoding SMILES and AA sequences, we use LASSO to select the best descriptors from various chemical fingerprints, physio-chemical properties of the compounds, and sequence-based and structural descriptors of the protein targets. We then employ sophisticated and flexible embedding methods for numerical descriptors [22], and pass the embedded input through several layers of adjusted transformer encoder layers to learn the descriptors. We further pay attention to the combined compound-target features to understand their intricate relationships. As a result, we produce accurate DTI predictions with the powerful attention mechanism without abandoning the diverse and informative compound-target descriptor sets, thus paving the way for increasingly multimodal DTI prediction approaches.

In addition to evaluating the prediction methods using train-test split testing, we computationally validated our methods on a huge DTI data set (185,676 bioactivities) from drug-target pairs provided by ChEMBL-V33 that were non-overlapping with the training dataset. Finally, after building the prediction methods, we tested those to predict novel (previously untested) bioactive interactions for 3492 approved drugs in ChEMBL-V33, opening new avenues for drug repurposing. As far as we know, no such comprehensive attention-based methods exist to seven superfamilies of the druggable targets, trained and validated on such big datasets as proposed in this study.

## 2. MATERIALS AND METHODS

### 2.1. Training dataset

The training data in this project consists of bioactivity values between drug-target pairs. We utilize two continuous label types for activity: DrugTargetProfiler (DTP) [23] scores and pChEMBL values. DTP scores are continuous values between 0 and 1, where 0 is considered inactive. DTP score is computed based on bioactivity values, assay format and target family, hence avoiding the shortcomings due to heterogeneity in activity values in public databases such as ChEMBL or BindingDB. Furthermore, the use of DTP scores facilitated our winning success in the CTD dream challenge [24]. On the other hand, pChEMBL value is defined as the negative log of molar concentrations in terms of IC50, EC50, XC50, AC50, Ki, Kd and potency. Higher pChEMBL (e.g. >6) represents a more active interaction, and lower means inactive (≤5).

We collected the training data for active targets from DTP, and verified inactive targets from PubChem using API, filtering out all non-human targets. All data points include both DTP scores and pChEMBL values as labels. The active dataset contained some duplicate drug-target pairs with differences in the recorded activity values, possibly due to different assay formats or detection methods. We handled this discrepancy by taking the median of the reported values. After median filtering, we removed any dubious pairs found in both the active and inactive datasets. We further removed those inactive pairs, where the DTP score was 0 but pChEMBL value was higher than 5, or vice versa. The resulting training dataset contained 947,195 interactions between 452,296 compounds and 1,251 targets. Within the compounds, 1,884 are approved drugs, 120 in phase I, 268 in phase II, 336 in phase III, and the remainder are preclinical compounds.

We divided the full dataset into seven parts according to the respective target superfamilies. Finally, we sub-sampled 20% of each set for testing and kept the remaining 80% for training (i.e., one-time train-test spit, instead of cross-validation).

### 2.2. Workflow of proposed method

Figure 1 presents the general workflow of our DTI prediction approach. After collecting and preprocessing the pairwise activity values from PubChem and DTP, we split the data into seven parts based on the respective protein superfamilies. We compute comprehensive, multimodal descriptor vectors composed of chemical fingerprints and physicochemical properties for the compounds, sequence-based descriptors, Zernike descriptors and subclass labels for the targets. Then, we robustly select the most informative descriptors with LASSO, separately for each of the seven protein superfamilies.

**Figure 1:**
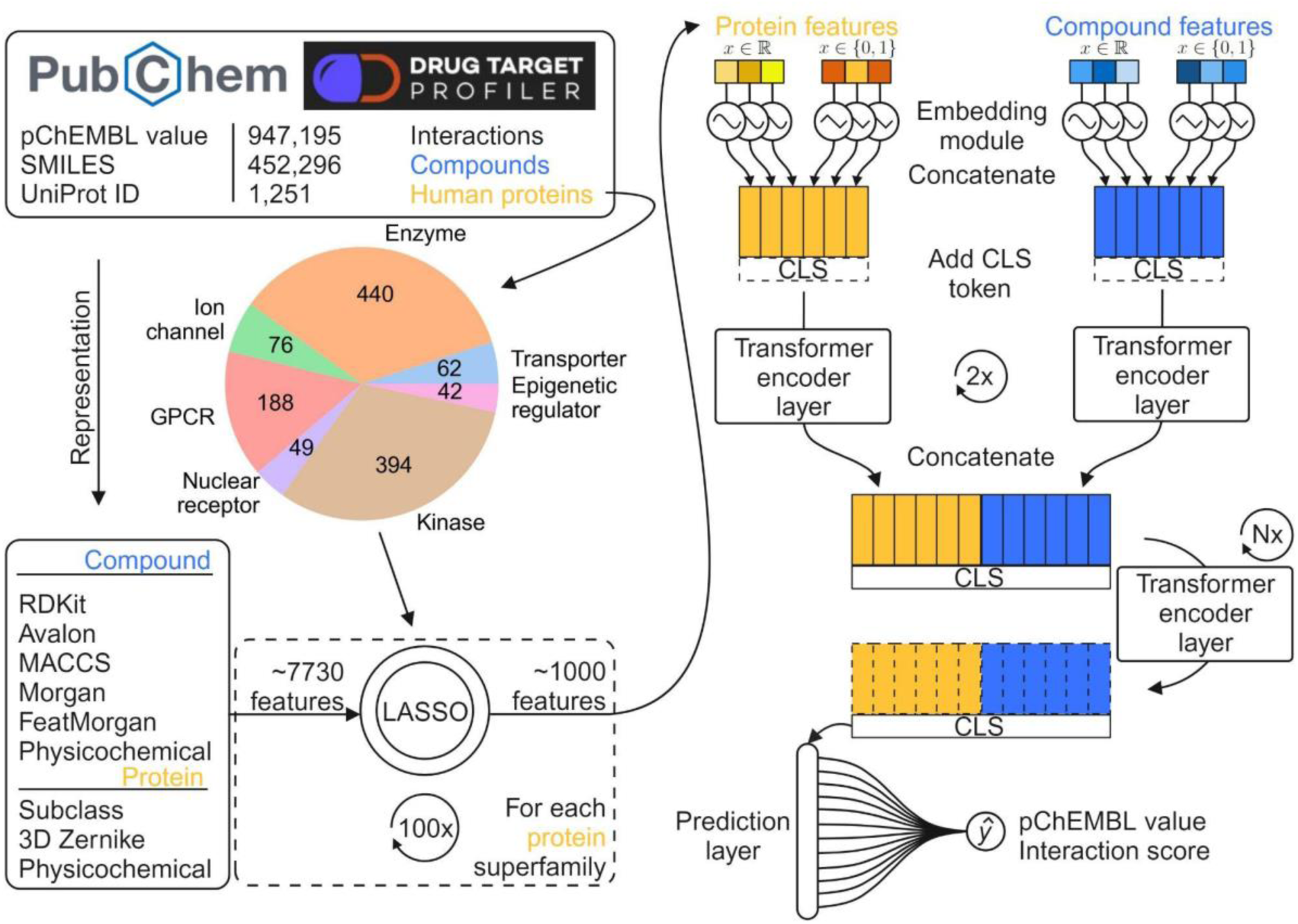
Workflow for the attention-based models to predict target bioactivities of approved drugs across seven protein superfamilies. Active interactions are obtained from Drug Target Profiler (DTP), and in-actives from PubChem. We generated a high-dimensional descriptor vector from the data to represent compound-protein pairs. We selected the most relevant features with LASSO, resulting in one feature set for each protein superfamily. We repeat the selection 100 times for robustness and retain the most frequently chosen descriptors. We separate each set into protein and compound descriptors and pass them into specialized embedding modules. The embeddings are then concatenated, appended with a CLS token, and fed into a series of transformer encoder layers. Finally, the transformed CLS tokens are passed into a prediction layer to yield a pairwise binding affinity prediction. Each protein superfamily will have two models, one to predict pChEMBL values and the other to predict DTP interaction scores (0-1).

With the prepared datasets, we trained 14 attention-based models: one for predicting pChEMBL values and the another for interaction scores, separately for the seven superfamilies. The architecture is uniform for all models. It consists of separate embedding modules for binary and continuous descriptors of both compounds and target proteins, followed by a series of transformer encoder layers. The encoder layers apply self-attention on the descriptor matrices equipped with CLS tokens and learn the compound-target descriptors and relationships. Finally, the CLS tokens are extracted and fed into a prediction layer that yields the final point prediction. The detailed model architecture is shown in **Supplementary Figure 1**.

### 2.3. Attention based architecture

We used advanced deep learning architecture for our prediction problem. The right side of Figure 1 shows the overall architecture, whereas **Supplementary Figure 1** illustrates the detailed architecture of the model. The model is a modification of the FT-Transformer [25]. The FT-transformer is an effort to improve the performance of neural networks when working with tabular data. It utilizes the encoder part of the transformer model [10] as its central learning tool, which relies heavily on self-attention. The distinct quality of the FT-Transformer over most other attention-based models is its ability to use numerical data in a tabular format as input instead of requiring data in text or image format. Therefore, our model is separate from using SMILES and AA sequences as central compounds and target descriptors while leveraging the self-attention process. Furthermore, our model design is not confined to the numerical descriptors used in this work but can be incorporated with any numerical descriptors. This flexibility allows for straightforward improvements in the future when superior compound or protein descriptors are discovered.

We group the input descriptors to our model into four vectors: continuous compound descriptors, binary compound descriptors, continuous protein descriptors, and binary protein descriptors, together describing one compound-protein pair. We first pass the vectors into an embedding module. The module converts an n-dimensional vector into a tensor of size n × d, where d is a tunable embedding dimension. While embedding is usually done for nonnumerical descriptors such as letters and words, we follow the unconventional manner of Gorishniy et al. [22] and use embeddings for numerical descriptors. We used a periodic activation function combined with a fully connected layer and an activation function for continuous descriptors. The embedding module for a continuous descriptor x is as given by the following equation:

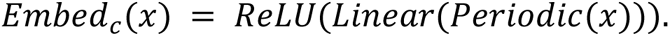

Here, ReLU and Linear refer to the Rectified Linear Unit and a fully connected layer, respectively. The periodic activation function (Periodic) converts a scalar input to a vector of arbitrary dimension. We define it as

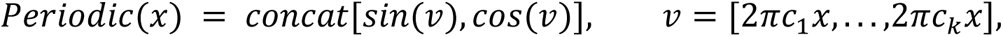

where *x* is the scalar input, *k* is an arbitrary activation dimension and *c*_*i*_ is a trainable parameter sampled from normal distribution *N*(0, *σ*), where the value of *σ* is determined by the practitioner. Thus, an input data vector *x* ∈ *R*^*n*^ can be converted via periodic activation into a matrix *A* ∈ *R*^*n*×2*k*^, therefore rendering it suitable for the attention mechanism. The embedding module is a simple lookup table with a cardinality of two without any additional layers for binary descriptors. All embedding modules are independent descriptor-wise, i.e., no weights are shared between embeddings of distinct descriptors.

After the embedding procedure, the model concatenates continuous and binary descriptors while keeping the compounds and proteins separate. A CLS token row of initially random values is also appended to the embedding tensor. The CLS token is originally introduced in the BERT architecture [13]. The model then passes the resulting tensors into a series of consecutive Transformer encoder layers. We do not utilize positional encoding in the encoders because the order of individual elements in descriptor sets does not matter. We apply self-attention to compound and protein descriptors separately during the first two layers, concatenating them into one tensor allowing the subsequent encoder layers to attend to all the features together.

Concerning the encoder layer structure, we closely follow the FT-Transformer and use the pre-norm variant of the encoder, where the residual connection branches out before layer normalization. This contrasts with the original Transformer, where residual branching occurs after normalization. The pre-norm variant facilitates the optimization process during model training [25]. Furthermore, we apply dropout to multi-head self-attention and fully connected layers in the encoder.

Following the encoder layers, we extract the CLS tokens and discard all the other rows in the data matrix. This is possible due to the encoder layers changing the values of the CLS tokens via the attention mechanism such that they gain sufficient knowledge and context about the compound-target descriptors. Thus, the CLS tokens contain enough information in a condensed form that they alone are passed into the prediction module for the final interaction strength point prediction:

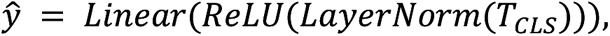

where *T*_*CLS*_ refers to the CLS token. We conducted the training on 4 Nvidia Volta V100 GPUs and 40 Intel Xeon Gold 6230 (2,1 GHz) CPUs. We parallelized the training with the PyTorch Distributed Data Parallel framework together with the Ray library functionalities. The training times for each model are recorded in **Supplementary Table 1**. We trained 14 predictive models, such that each of the seven protein superfamilies had two separate models for pChEMBL values and DTP scores. We initialized the weights of linear layers with Kaiming initialization. For loss calculation, we used the logarithmic hyperbolic cosine loss function. The function is known to be less sensitive to outliers and is defined as given the following equation:

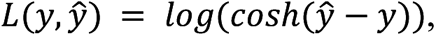

where *y* is the true label, and *ŷ* is the prediction. During preliminary testing, we observed that the log-cosh loss yielded better results than the commonly used mean squared error. For optimization, we opted for the AdamW optimizer.

We set the limit for training to a maximum of 500 epochs and utilized early stopping with a patience of 10 epochs. Patience refers to a period after which the training terminates if the test loss does not decrease. This is done to avoid overfitting and to preserve model generalizability. We checkpointed the model weights after each epoch and retained five checkpoints with the lowest reported test losses. We used the checkpoints as an ensemble for later predictions and inference.

### 2.4. Descriptor sets

Instead of relying on text-based descriptors generated by attention-based models, we analyzed various comprehensive sets of numerical descriptors ever reported in drug-target-based prediction methods in the literature. Some descriptor sets are related to compounds, whereas the remaining are associated with target proteins. The following sections describe different descriptor sets adapted in the proposed method.

#### 2.4.1. Compound descriptors

All compound descriptors in the proposed study take as input the canonical Simplified Molecular Input Line Entry System (SMILES) representation for a compound. SMILES is a widely used system that notates the chemical structure of a compound in linear string format. We use the SMILES to calculate several fingerprints and physicochemical descriptors for the compounds in training and test datasets.

##### 2.4.1.1. Chemical fingerprints

Chemical fingerprinting is a method to capture the identity of a molecule in a fixed-length, binary format based on different properties ranging from shape and connectivity to chemical descriptors associated with molecular interactions. For binary fingerprints, each bit represents an atom’s presence (1) or absence (0). The length of the bit vector varies depending on the fingerprint used. Fingerprints are used to compare or analyze the structural similarities between the compounds, termed quantitative structure-activity relationship (QSAR). Structurally similar compounds may often be assumed to have similar activity towards a target [26]. Tanimoto or Jaccard coefficient (*T*_*c*_) is a standard metric to compute structural similarity between fingerprints of two compounds as given by the following equation:

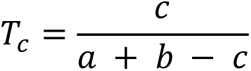

where a and b are the number of atoms present in each compound fingerprint, and c is the intersection of atoms between the two compounds. The *T*_*c*_contains real values between 0-1, where a higher value corresponds to higher structural similarity between two compounds. Importantly, different fingerprint types will produce different similarity scores. This may result in so-called activity cliffs, where similar compounds exhibit different activity towards a target, thus going against the QSAR above assumption. To overcome this, studies have found it beneficial to incorporate several fingerprints for a more robust view instead of relying on only one scheme [27],[28]. We, therefore, adapted five different fingerprints in our work: Molecular Access System (MACCS) keys [29], RDKit [30], Avalon [31], Morgan [32], and Featmorgan [32] fingerprints. MACCS keys are 166-bit-long vectors, where each bit corresponds to the presence of pre-specified structural fragments. RDKit fingerprint is a path-based algorithm where molecule fragments are located by following a molecular bond path, after which they are mapped to a fixed-size bit vector with a hash function. Avalon fingerprint uses a similar mechanism but considers slightly different molecule descriptors in encoding. Morgan fingerprint similarly uses a hash function but identifies the fragments by evaluating their atomic environment within a circular radius instead of a linear path. Finally, as a variation of the Morgan, Featmorgan also uses the circular approach but encodes different chemical properties of surrounding atoms. We used a radius of two for the Morgan and Featmorgan fingerprints, rendering them analogous to ECFP4 and FCFP4, respectively. The RDKit, Morgan and Featmorgan fingerprints used in this work are each 2048 bits long, while the Avalon fingerprint is 512 bits.

##### 2.4.1.2. Physicochemical descriptors for compounds

We used physicochemical properties to complement the molecular fingerprints in compound descriptors. The physicochemical properties of a compound refer to the physical and chemical characteristics that determine its behavior in different environments. They have a direct influence on how compounds interact with target proteins. In this work, we use RDKit to calculate a vector of 200 continuous, real-valued descriptors, each denoting the magnitude of a compound physicochemical property. Some examples of physiochemical descriptors used in this study are related to size, surface properties, partial charge, molecule fragmentation, drug-likeness, and many others.

#### 2.4.2. Descriptors for target proteins

Computing numerical descriptors for protein targets is a complex task with various existing methodologies [33]. Descriptors based on protein 3D structure have become an attractive approach due to the ever-increasing data on 3D protein structure, complemented by the accurate predictions of AlphaFold [16]. A recent protein shape retrieval challenge [34] compared the performance of several such descriptors. A method based on 3D Zernike moments [35] was consistently among the top performers. We adopted this approach to represent the 3D structure-based descriptors for the protein surface as a numerical vector. We obtained the protein 3D structures as Protein Data Bank (PDB) files to compute the Zernike descriptors. We collected the experimentally validated structures for 1141 proteins with UniProt ID search. We used AlphaFold high-confidence (>90) predictions for the remaining 270 proteins in this study. While it is known that for certain protein classes, such as kinases, using the binding pocket structure instead of the whole protein yields more accurate results [36], we used the whole protein structure for more direct comparison and uniformity across the protein superfamilies. In addition to the Zernike descriptors, we used sequence-based descriptors to exploit patterns between individual proteins and further classified the proteins into subclasses.

##### 2.4.2.1. Zernike descriptors

Zernike descriptors are a series expansion of a protein surface as a 3D function. The surface function is of the spherical coordinate system form f(r, θ, ϕ), where ‘r’ is the radius, ‘θ’ is the polar angle, and ‘ϕ’ is the azimuthal angle. A protein is voxelized to obtain the functional form of the surface. In the voxelization procedure, the protein is placed on a 3D grid based on the coordinates of its amino acids. Each atom is approximated by a Gaussian and integrated to obtain voxel points. Now, we can express the surface function as

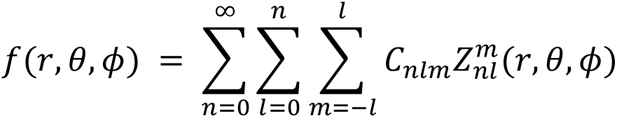

where *C*_*nlm*_ are Zernike moment coefficients and 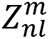 are 3D Zernike polynomials with order ‘n’, degree ‘l’ and repetition ‘m’. The polynomials are basis functions, defined as

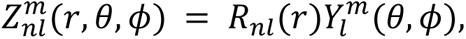

where 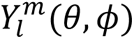 are spherical harmonics and *R*_*nl*_(*r*) is a radial function:

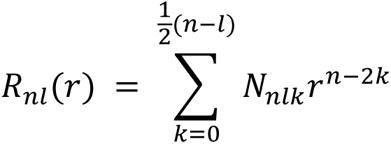

where N is a normalization factor. We find that the spherical harmonics ‘Y’ depend on the angles ‘θ’ and ‘ϕ’, while the radial function R only depends on the radius. Let us transform the function ‘f’ to the Cartesian system: f(r, θ, ϕ) → f(x). As orthonormal basis functions, the series expansion of the expression is given by

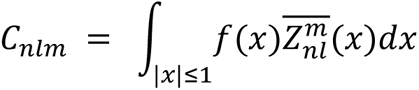

Finally, we take the norm to obtain the complete Zernike descriptors:

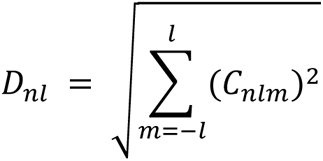

These descriptors are rotationally invariant, meaning there is no need for pre-alignment of proteins before calculating the descriptors. Furthermore, the resolution of the surface representation, i.e., the dimension of the descriptor vector, can be adjusted by altering the value of the order ‘n’. We based our value of n on literature [37] and set it to 20 to yield a 121-dimensional numerical vector. For our convenience, the Python code for calculating the descriptors was supplied by Di Rienzo et al. [38], who used these successfully in their research.

##### 2.4.2.2. AA sequence-based descriptors

To complement the 3D-based Zernike descriptors, we include sequence-based descriptors for proteins (576-dimensional feature vector). These descriptors are related to mono-peptide and bi-peptide amino acid frequency compositions, polarizability, hydrophobicity, and aromaticity. Our GitHub repository contains scripts to compute all these descriptors.

##### 2.4.2.3. Protein subfamily information

We used binary labels to further classify proteins in each superfamily e.g. Tyrosine kinases is a subfamily of Kinase, Chemokine and Amine are the sub family of GPCRs. This protein subfamily information is extracted from the ChEMBL database.

### 2.5. Feature selection using LASSO

By combining descriptors from the nine descriptor sets, we obtained 7730 descriptors in total. Such a huge descriptor vector consumed time to train the prediction algorithm on a dataset with ∼1M interactions. To speed up the training process, we performed feature selection to reduce the dimensions of the compound-target descriptor sets. We separately selected the descriptors for each target superfamily using the same protocol. With an adequate selection mechanism, only the most important descriptors are retained, while less significant ones are discarded. Crucially for our work, feature selection allows for faster training times and lower system memory requirements. Often, it also improves the signal-to-noise ratio of data and counters overfitting [39]. We adapted the least absolute shrinkage and selection operator (LASSO). Using LASSO for selecting descriptors is straightforward as it reduces the coefficients of less important variables to zero. Therefore, there is no need to set any arbitrary thresholds for selecting descriptors, as would be the case with, for example, ridge regression. LASSO also maintains interpretability by not altering the original form of the descriptors, unlike methods such as principal component analysis. Recalling our data format of (*xi, yi*), i = *1, 2, …, N,* the minimization objective of LASSO is defined as given by the following equation:

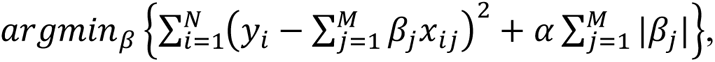

where α is a constant penalty term and (β1, …, βj) are the coefficients to estimate. We used a separate subset of data for each protein superfamily to solve the optimization problem. In the subsets, we included all interactions involving approved drugs and phase III investigational compounds with respective protein superfamilies within our dataset. Using subsets allowed for easier and faster convergence than using the full dataset. Protein subfamily labels were excluded from the selection process and were thus always included in the final feature vector. We determined the value for α using an iterative fitting scheme along a regularization path with 5-fold cross-validation. The cross-validation was performed separately for each protein superfamily on the data subsets. After finding the best α, we fit the model once again with the full subset and kept the features for which β > 0. Repeating the procedure on the same subset often resulted in a different α and consequently a different number and combination of selected features. This occurred most likely due to variance in the data splitting within separate cross-validation initiations. Thus, for more robust results we conducted stability selection, i.e., we ran the LASSO selection procedure 100 times and retained the most frequently selected features. The subset of data was kept the same at each instance, only the data folding was randomized. As a result, we obtained seven sets of features, one for each protein superfamily.

Figure 2 shows the superfamily-wise representation of selected descriptor sets. We fixed the number of descriptors to select 1000 (excluding protein subclass labels), thus resulting in an approximately eight-fold decrease in descriptor counts. Zernike and other protein-based descriptors were among the most important selected descriptors. We make further analysis of descriptor set usage in Figure 3. Interestingly, nearly half of the compound descriptors are not used in any protein superfamily. For instance, none of the compound descriptors are general enough to be used in all 7 superfamilies. In comparison, the protein descriptors are used much more frequently.

**Figure 2:**
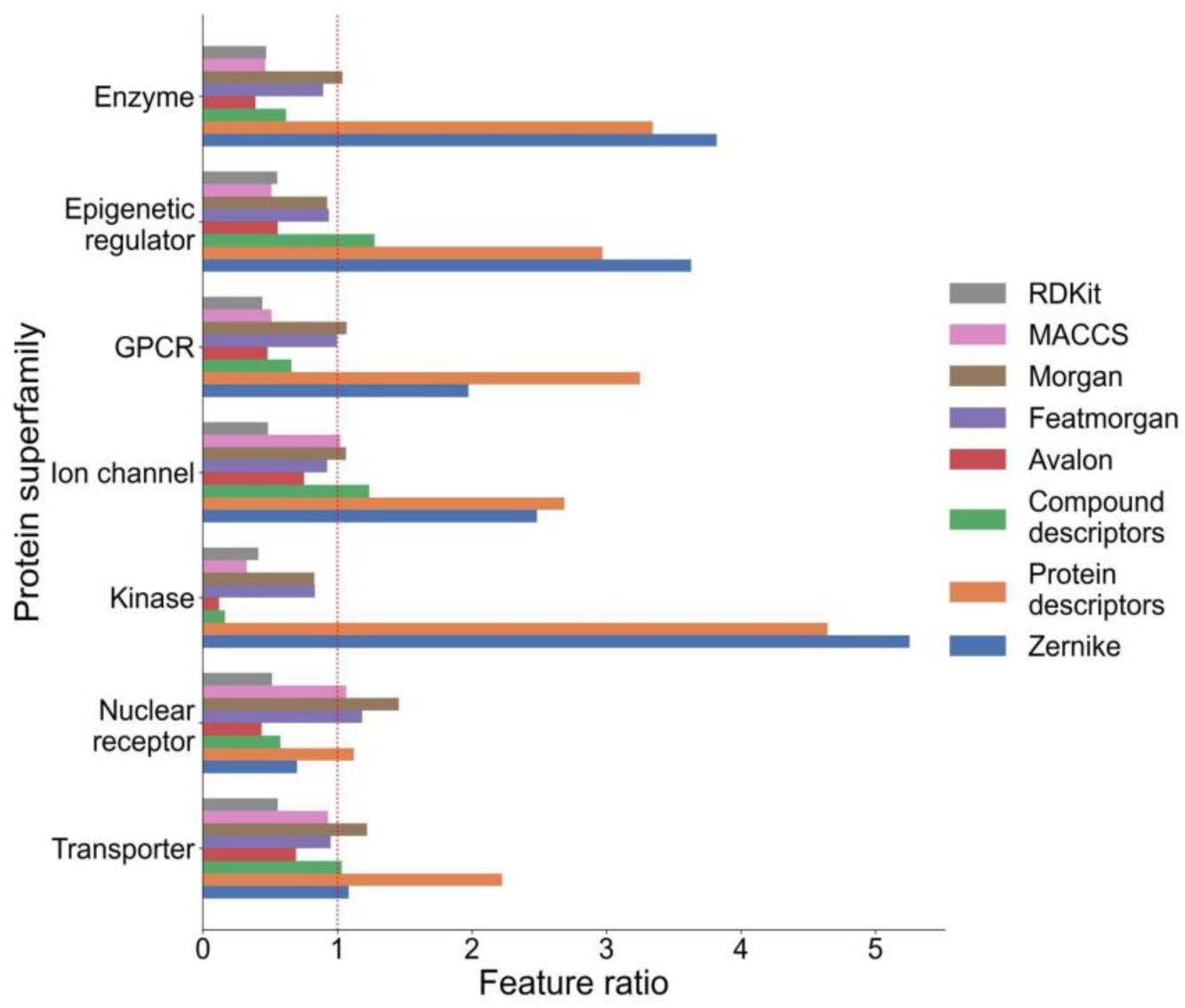
The change in descriptor vector dimension after feature selection. Originally, each compound-protein pair was represented by a 7719-dimensional vector. We reduced the dimension to 1000 with LASSO. Note that the selected descriptors are different across protein superfamilies. The horizontal bars show the direction of change in the dimension of separate descriptor sets to the 1000-dimensional descriptor vector for each superfamily when compared to the original, 7719-dimensional vector. Bars that cross the dotted reference line indicate an increased contribution of a descriptor set to the overall composition of a feature-selected vector than to the original vector. We omit protein subfamily labels from the comparison as they are always included in the descriptor vectors.

**Figure 3:**
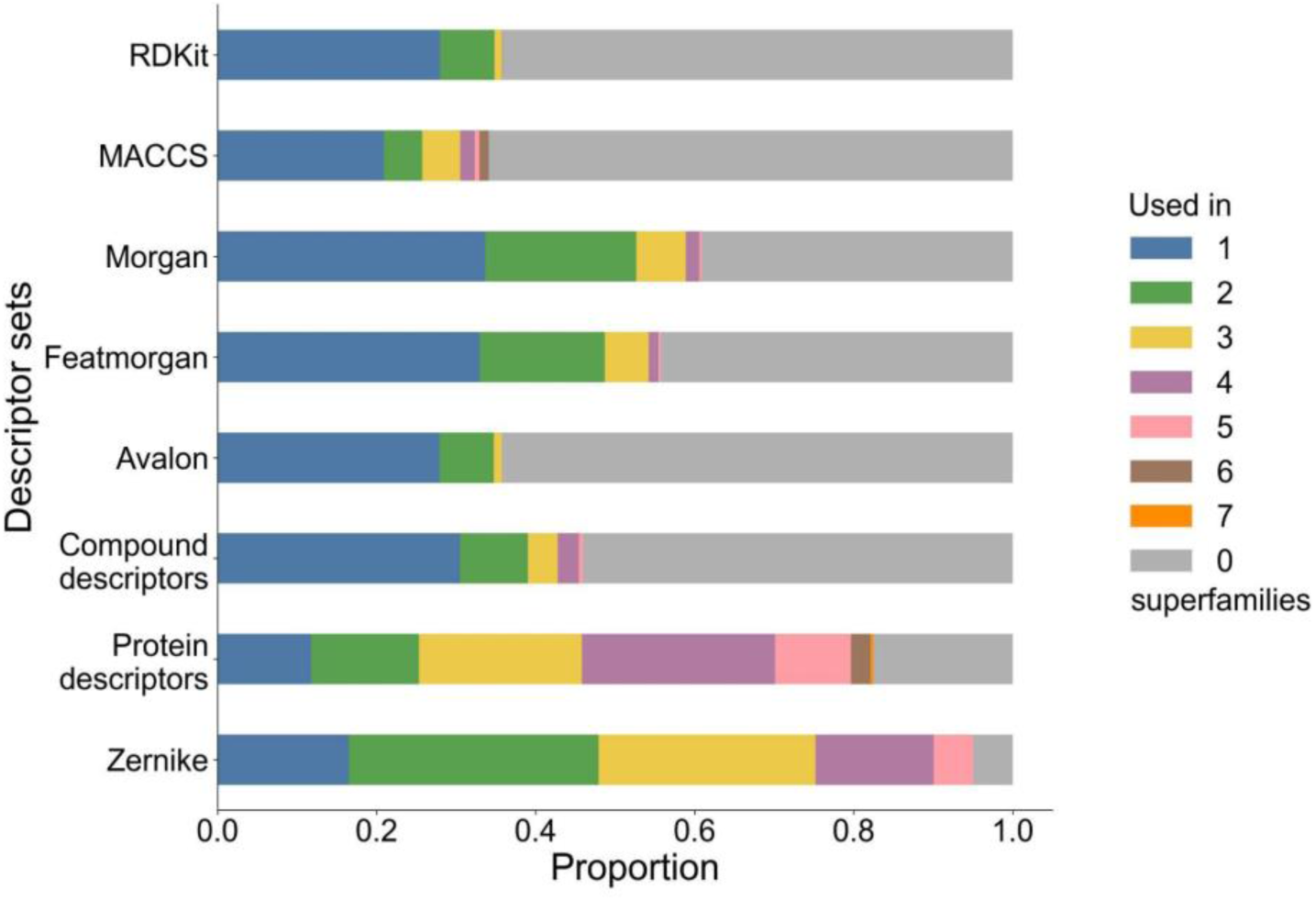
Proportions of how often individual descriptors belonging to a descriptor set are selected by LASSO into a model for each protein superfamily. The descriptors are grouped by their generality, i.e., how often they are selected for different superfamilies. The proportion of descriptors selected for all seven superfamilies is the lowest, while the proportion not contributing to any superfamily is the highest.

### 2.6. Hyperparameter tuning

The hyperparameter optimization technique used in this work was Heteroscedastic Evolutionary Bayesian Optimization (HEBO) [40]. We chose this method because it won the competitive NeurIPS black-box optimization challenge in 2020 [41]. HEBO improves on two weaknesses found in plain BO approaches. Firstly, BO often assumes neat, Gaussian noise likelihoods. Instead, HEBO considers data heteroscedasticity, meaning that it does not assume constant variance in the data. This is achieved with non-linear input and output transformations. Secondly, BO usually utilizes one acquisition function that is assumed to be optimal. However, different acquisition functions can easily lead to conflicting results. Thus, HEBO uses an algorithm called multi-objective acquisition ensemble to search for the best solution via a Pareto front [42]. In our implementation, we additionally combine HEBO with Hyperband [43] for reduced computational cost. Hyperband accelerates the optimization process by shutting down less promising runs early on.

We utilize a simple train-test split in our hyperparameter optimization approach. We selected this method instead of the more robust cross-validation approaches due to constraints on time and computational resources. 80% of the original data was used for training and 20% for testing. Thus, for each superfamily separately, we train a model with initially random hyperparameters on the train split. The resulting model is evaluated on the test split. Based on the evaluation, HEBO selects new hyperparameters, and the cycle is repeated until the budget limit is reached. We then select the best hyperparameters based on the evaluation metrics gained from the test set. Finally, we test the model on separately collected, independent test data that is fully excluded from the optimization procedure.

**Supplementary Table 2** lists the range of search spaces for each hyperparameter, as well as the constant hyperparameters. We set the budget for tuning at 144 hours or 50 training iterations for each class, depending on which limit was reached first. We set the Hyperband grace period to 5 epochs, meaning that no iteration would be terminated before then. For a fair comparison, we kept the search spaces the same for all proteins, with the exception of the number of training epochs. We varied it between 50 and 75 depending on the observed speed of convergence and the size of the training set of a protein superfamily. **Supplementary Table 3** shows the best resulting hyperparameter configurations for all the models.

### 2.7. Performance metrics

We evaluate our models with three metrics commonly used in DTI regression problems: root mean squared error (RMSE), Spearman’s rank correlation coefficient, and concordance index (CI). Each metric gives a distinct insight to the model performance. Thus, it is important to consider them together as a whole, instead of focusing on only one stand-alone metric. RMSE measures the magnitude of error between predicted and actual activities, defined as

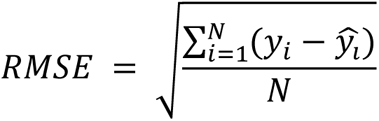

where y is the true label, *ŷ* is the prediction and N is the total number of datapoints tested. A lower RMSE value indicates a better fit of the model to the data.

Spearman correlation describes how well the relationship between the predicted and true labels can be described using a monotonic function, i.e., if the predicted values increase or decrease in tandem with the labels. It does not assume a linear relationship between the variables and is thus less sensitive to outliers. The correlation produces values between −1 and 1, where −1 indicates perfect inverse monotonic, 1 perfect monotonic, and 0 no monotonic relationship. Spearman correlation is defined by the following equation:

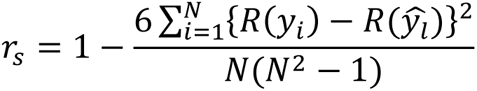

where R(·) is the rank of a variable. In the context of our study, higher Spearman correlation is better.

In the context of DTI prediction, CI refers to the likelihood that, when randomly selecting two drug-target pairs, the predictions made on those pairs are in the correct order in terms of label value. CI is defined using the following equations:

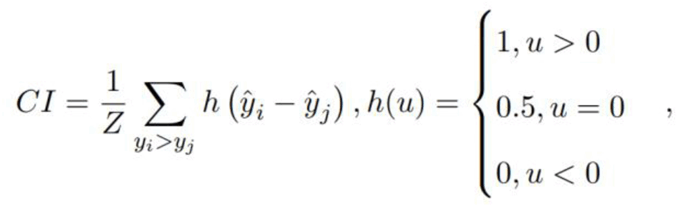

where Z is the number of distinct label values in the data. An ideal model achieves a CI of 1.0, while predictions obtained by random guesses will yield a CI of 0.5.

## 3. RESULTS AND DISCUSSIONS

### 3.1. Model training and testing

We assessed the performance of the proposed methods with a testing dataset separated from the original data prior to training. The train-test split was obtained by randomly sampling 20% of the full data for testing and hyperparameter tuning, while the remaining 80% was used for training. The approach is more resource-efficient compared to other common techniques, such as k-fold cross-validation. The random sampling resulted in near-identical bioactivity distributions between training and testing sets for each superfamily? (**Supplementary Figures 2 and 3**), due to the large size of the original, unsampled data set. We conducted the data-splitting process for each protein superfamily separately.

Figure 4 shows the testing results and contour plots for predicting pChEMBL values and interaction scores. The results are based on a mean of predictions from five model checkpoints of different weights. The checkpoints are obtained by saving model weights after each epoch during a training run, and then selecting those with lowest reported test losses for use in inference and prediction. Such an ensemble approach slightly improved the performance without adding overhead to the training. The prediction times do increase with the ensemble but remain sufficiently low. We also reported average prediction time for an interaction across each of the seven superfamilies in **Table 1**. The times vary mainly due to differences in the model sizes.

**Figure 4:**
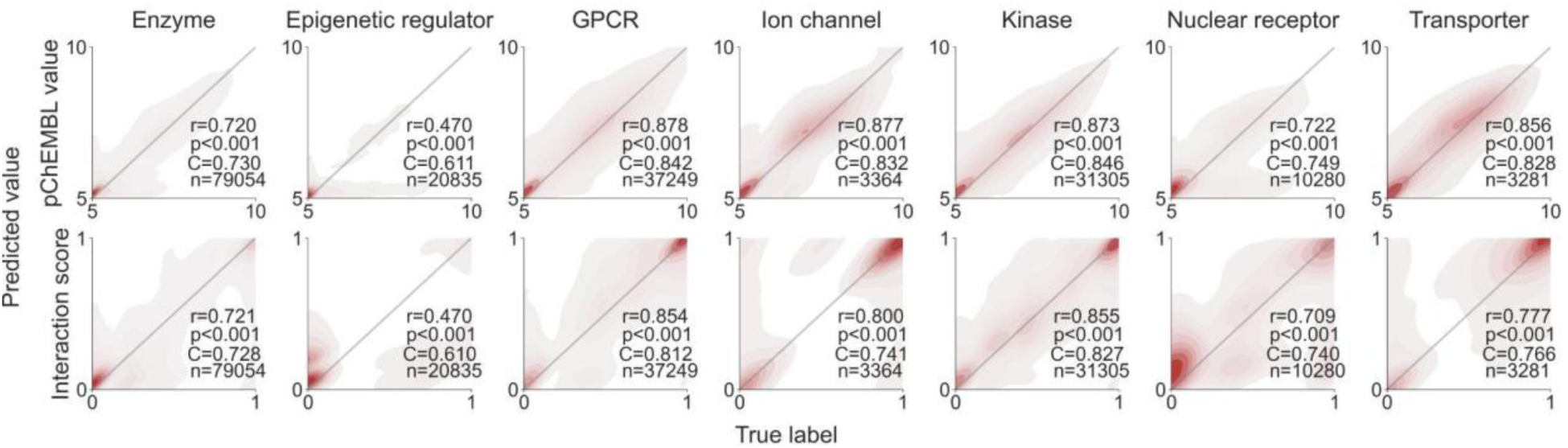
Contour plots of the testing results for each protein superfamily. The top and bottom rows correspond to models predicting pChEMBL values and interaction scores, respectively. The gray diagonals represent the ideal results, where predictions match perfectly with the true labels. Each plot is annotated with the Spearman correlation coefficient (r), the associated p-value (p), the concordance index (C), and the number of interactions (n).

**Table 1:** Testing results from final training after hyperparameter tuning. The results are derived from an ensemble of five best checkpoints with the best test results in each training run. We report the time taken to predict one drug-target interaction with each ensemble in milliseconds.

| Protein | Spearman | RMSE | CI | Prediction time<br>for one pair (ms) |
| --- | --- | --- | --- | --- |
| Enzyme | 0.720 | 0.509 | 0.866 | 34.6 |
| Epigenetic regulator | 0.470 | 0.560 | 0.811 | 16.1 |
| GPCR | 0.878 | 0.679 | 0.865 | 34.2 |
| Ion channel | 0.877 | 0.644 | 0.875 | 32.9 |
| Kinase | 0.873 | 0.625 | 0.861 | 37.7 |
| Nuclear receptor | 0.722 | 0.779 | 0.822 | 9.1 |
| Transporter | 0.856 | 0.696 | 0.848 | 35.2 |

The test results imply that the model design leads to strong learning capabilities across protein superfamilies. The exception here is the model for epigenetic regulators. By the tail of the contour plot in Figure 4, we notice how the epigenetic regulator model tends to underestimate the activity of compounds, often predicting them as inactive. A similar trend is visible in the nuclear receptor model and, to some extent, in the enzyme model. We attribute this behavior to the skewness of the training data label distribution towards inactive values (as shown in **Supplementary Figure 2**). For the other models, the predictions align well with the true labels.

### 3.2. Model validation on independent test set

The testing results alone are insufficient to evaluate model performance. Testing reveals a model’s general behavior and learning capacity, but the results are often too optimistic. This is partly because the test set involves hyperparameter tuning, meaning that the model is optimized to make predictions on the training dataset. This issue is often solved by splitting the test set into two sections, one for tuning and the other for testing. Even with this technique, the prediction setting is too easy for a practical performance evaluation because the compounds and proteins in the test set likely overlap or are similar to those used in the training data.

To obtain a more objective understanding of our models’ robustness and usefulness, we collected a new, independent validation dataset. The validation set includes those compound-target pairs present in ChEMBL-V33 reported in articles/documents not considered in the train-test split sets. Similar to the training data, some drug-target pairs contained multiple conflicting entries. We retained the median label value of such cases and discarded the rest. Furthermore, we removed interactions for some proteins for which there were no Zernike descriptors available. The resulting validation set consists of 185,676 interactions between 119,034 compounds and 1,681 targets. This independent validation set is provided in **Supplementary File 1** to help reproduce our comparative analysis.

With this external validation set, we designed three validation scenarios of increasing difficulty, as shown in Figure 5. The first and easiest case is bioactivity imputation, where the validation set contains compounds and proteins already present in the training set; only the compound-target pairings are different. The second scenario includes interactions between unseen compounds and seen proteins. Finally, the third and most challenging scenario contains only compounds and proteins not seen in the training set. We omitted the apparent fourth case of known compounds and unknown proteins due to inadequate data collection.

**Figure 5:**
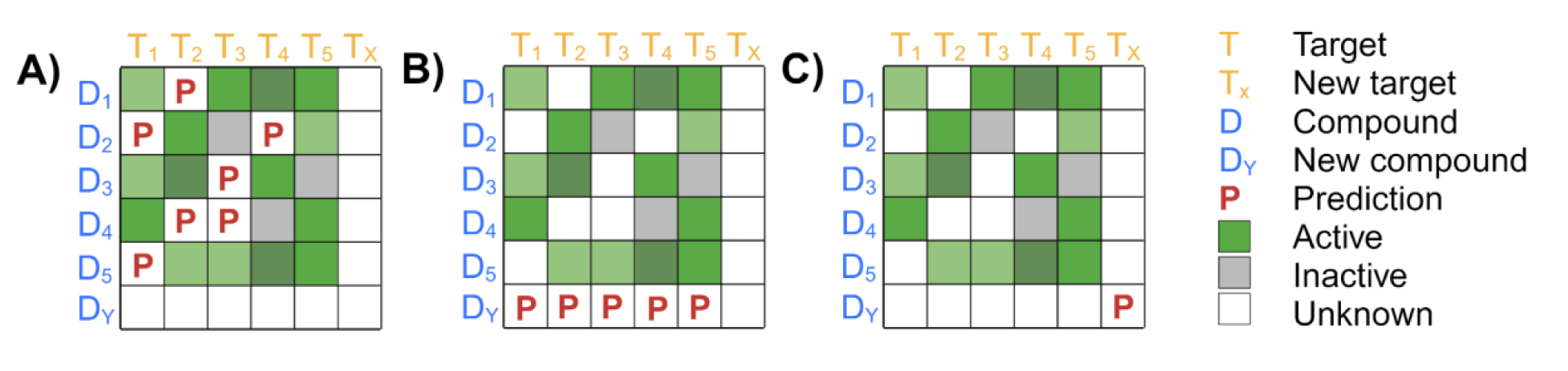
An illustration of the three validation scenarios. A): bioactivity imputation, where all validated compounds and targets are available in the training set, B): new compounds, where none of the compounds but all targets are available in the training set, C): new compounds and new targets, where neither the compounds or proteins are available in the training set.

We evaluated the performance in the three independent validation scenarios across seven superfamilies. Similarly to the testing results, the results are from an ensemble of five best checkpoints. Figure 6 displays the contour plots, the Spearman correlations and their associated p-values, the number of pairwise interactions, and the average Tanimoto coefficients (TC) for each protein superfamily and validation scenario. Spearman correlation and RMSE are also reported in **Supplementary Table 4**. We calculated the average TC by applying the similarity equation to find the similarity between all compounds in the training and validation sets and taking the mean of the resulting similarities. We used RDKit fingerprints for molecule representation, noting that different fingerprints would likely result in slightly different similarity scores.

**Figure 6:**
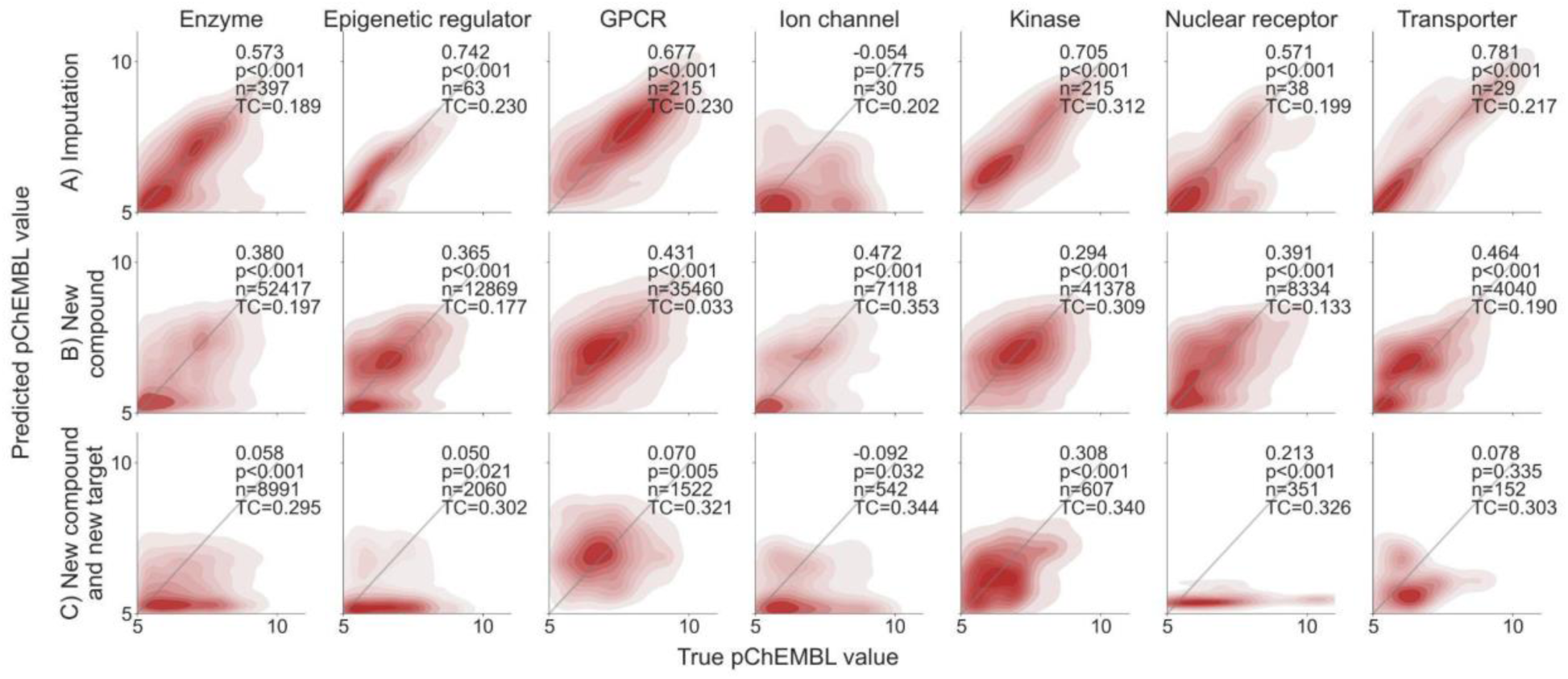
Contour plots of the validation results for each protein superfamily in three scenarios. The results are based on the pChEMBL labels. The gray diagonals represent the ideal scenario, where predictions match perfectly with the true labels. Each plot is annotated with the Spearman correlation coefficient and the associated p-value, the number of data points (n) and the average Tanimoto similarity (TC).

We observed unexpectedly low TC in the bioactivity imputation validation scenario. To address the seeming discrepancy of the simultaneous occurrence of shared compounds and low structural similarity, we note that the number of compounds is relatively low in the bioactivity imputation data sets. Thus, the existence of the same compounds between the compared sets has only a little influence on the overall similarity score of the sets. In general, compound similarities are low across validation sets (TC<0.35), thus increasing the difficulty of predictions in the validation set.

As expected, the performance generally decreases as the test cases become more challenging. The models generally perform well in bioactivity imputation, except for the ion channel model. Moreover, the epigenetic regulator performed surprisingly well in comparison to the train-test results shown in Figure 4. We note that the low number of available data points in the bioactivity imputation scenario leaves room for variations in results due to chance. This differs from the new compound validation scenario, where the datasets are sufficiently large. Despite the decrease in model performance, the overall results in this second scenario are satisfactory. We confirm an adequate predictive power across protein superfamilies when finding new compounds for known targets.

Interestingly, the only exception is the kinase model, where the performance surpasses that of the new compound’s validation scenario. We hypothesize that the inherent similarity between kinase structures diminishes the effect of unknown proteins, meaning that the last two scenarios are of similar difficulty for the kinases. For the other models, predicting the interaction between an unknown compound and an unknown protein is more difficult. This is a common phenomenon in contemporary DTI prediction methods, as this validation scenario is practically difficult for current methodologies and is mostly ignored in existing studies [44].

### 3.3. Comparison with other methods

Davis [45] provides large-scale benchmarking data from high throughput target activity screening, where each of the 442 human kinases is screened against each of 72 inhibitors (442×72=31824 interactions). While the Davis dataset is widely used in DTI research, its exact use in model evaluation differs between studies. For the sake of fair comparison, we closely follow the testing pipeline devised by Monteiro et al., the authors of DTITR [17]. Released in 2022, the DTITR method is relatively similar to ours, with the most significant difference being their use of encoded AA sequences and SMILES as input instead of curated features.

Monteiro et al. filtered the Davis data by only including proteins with residue lengths between 264 and 1400 and compounds with SMILES string lengths between 38 and 72 characters, resulting in 423 protein targets and 69 compounds. They then split the data into six folds, five used for training and hyperparameter tuning. The sixth fold was used for testing and comparison against other models. We utilized the same data and train-test folds, except for the further exclusion of 8 proteins for which Zernike descriptors could not be calculated and the removal of a few duplicates. We used the same three-dimensional structure for proteins with point mutations as for the corresponding wild-type proteins. This is because we were unable to find structures for mutant proteins. Subsequently, we repeated the training workflow for the kinase model based on the provided training folds, including feature computation and selection, hyperparameter tuning, and training. With near-identical training and testing data between models, the results can be attributed solely to differences in modeling approaches.

Figure 7 displays the results obtained from model predictions on the test fold (see details in **Supplementary Table 5**). In addition to our model and DTITR, the tested models are KronRLS [46], GraphDTA and some of its variants [47], SimBoost [48], Sim-CNN-DTA [49], DeepDTA [50], and DeepCDA [51]. All these comparison models have been trained on the same data by Monteiro et al. for consistency. Likewise, the reported results arise from predictions on the same test fold. Our model performs similar to DTITR, while outperforming the other models in terms of Spearman correlation and concordance index. DTITR does maintain a slight lead in terms of RMSE, with our model achieving the second-best performance together with DeepCDA. Overall, we reached a level of performance comparable to the current state-of-the-art.

**Figure 7:**
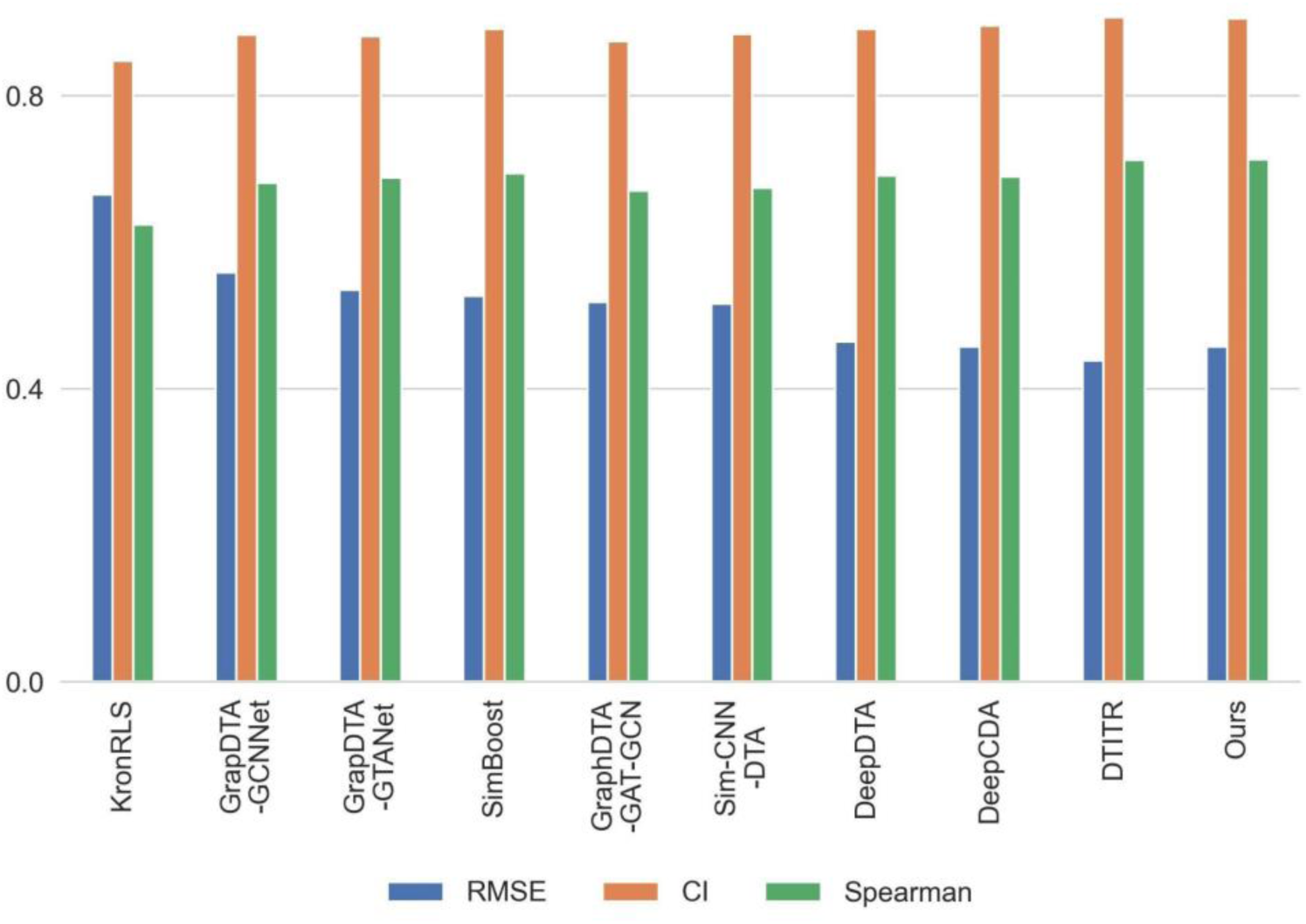
State-of-the-art comparison on the Davis dataset with three performance metrics. The results are derived from predictions on the test split constructed by authors of the DTITR method (more details in **Supplementary Table 5**).

### 3.4. Predicting novel bioactive interactions for approved drugs to open new avenues for drug repurposing

After successfully evaluating the robustness of the proposed prediction models using testing and validation datasets, we used the deployed models for each of the seven protein superfamilies to predict new and previously untested bioactive interactions for 3492 approved drugs against 1411 human targets. Figure 8 shows the distribution of known and predicted interactions across seven protein superfamilies. More details on the predicted and known interactions in terms of pChEMBL values are available in **Supplementary File 2**. Prediction of secondary active and inactive targets for approved drugs can lead to new drug repurposing applications and may become beneficial for finding new treatments for various diseases.

**Figure 8:**
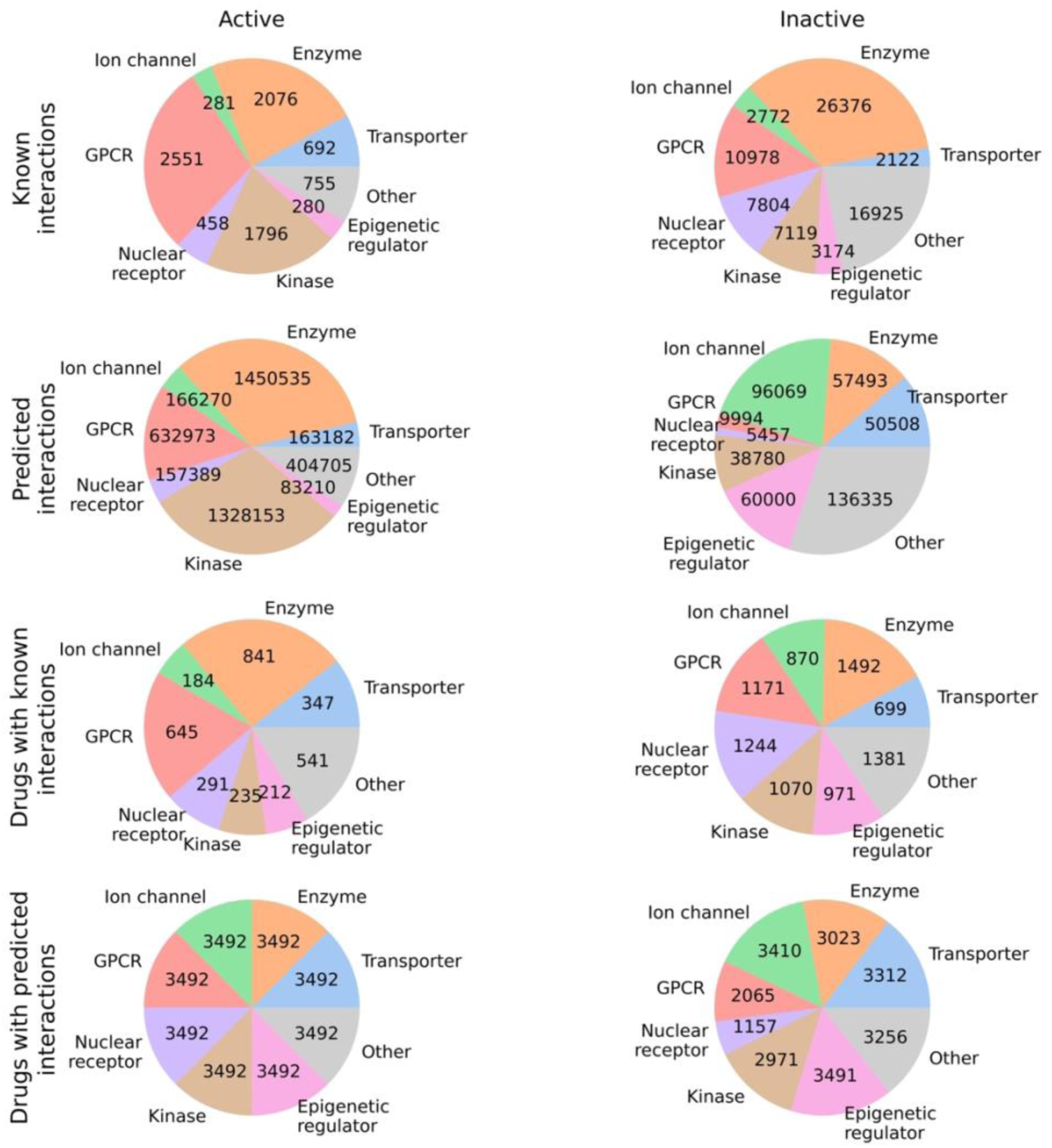
The number of known and predicted interactions and number of approved drugs for each protein class. The proportions are separated into active and inactive categories. An interaction with pChEMBL value of 5.0 or smaller is considered as inactive. For the predicted values, we set the inactivity threshold at 5.1 due to the regression model predicting many values close to, but not exactly, 5.0. **Supplementary File 2** shows more details on the predicted and known interactions of approved drugs across 1411 human proteins.

## 4. CONCLUSIONS AND FUTURE DIRECTIONS

In this study, we proposed attention-based methods to predict novel DTIs across seven superfamilies of the protein targets. Attention-based methods are the latest extension of deep learning and are proven to perform better than conventional machine learning or deep learning methods. Except for epigenetic regulators, prediction models for the remaining of the six super families showed high Spearman correlation >0.7. Low performance for targets from epigenetic regulators superfamily is most likely due to the skewing of the training data distribution towards inactive values. We also compared our proposed method with nine deep-learning and graph neural networks based on the latest methods reported in the literature for DTI prediction on the Davis kinase activity dataset, showing state-of-the art performance. We have provided the benchmark training dataset of 185,676 interactions across 119,034 compounds and 1,681 human targets. This comprehensive benchmark dataset (**Supplementary File 1**) can be utilized to develop and test future drug-target prediction methods.

Importantly, we also performed a comprehensive computational validation on the non-overlapping Drug-target pairs from ChEMBL-V33. This comprehensive dataset contains 185,676 interactions, 119,034 compounds and 1,681 targets that are not available in the PubMed training dataset. We defined three case scenarios to assess the validation results, as shown in Figure 5. Except for Ion channels, prediction models for most of the protein superfamilies performed well for two of the scenarios, i.e. missing value imputation (Spearman correlation > 0.57) and new unseen compounds (Spearman correlations > 0.36). Even for the third and most challenging scenario (new unseen compound and targets), kinase prediction models showed Spearman correlation > 0.3. This third scenario is not usually reported in DTI prediction studies. With such a high performance in testing and validation, the proposed prediction methods are expected to be generalizable to predict interactions with new compounds across seven protein superfamilies.

The proposed methods should especially be helpful during the hit identification phase in target-based drug discovery. We have provided all the training, testing and validation datasets, source codes, and user descriptions on GitHub (https://github.com/AronSchulman/MMAtt-DTA), so that researchers can easily reuse our methods or extend our analysis. Finally, we utilized the deployed methods for 1411 human proteins across the seven superfamilies to predict novel and previously unexplored interactions across 3492 approved drugs (with max phase of 4 in ChEMBL-V33). We completed the DTIs matrix for 3492 x 1411 with the help of known and predicted pChEMBL values (Figure 8**, Supplementary File 2**). Such a complete DTI matrix for approved drugs can open new avenues of drug repurposing and help understand their mechanism of action. Furthermore, such a complete drug-target matrix can further help improve the accuracy of in-silico drug combination prediction algorithms.

In future, we will explore language-based embeddings using SMILES and AA strings to investigate whether those can further improve the proposed approach. Some of the larger protein superfamilies can be divided into smaller groups, e.g., enzymes could be further divided into transferases and cytochromes. Similarly, some new protein superfamilies can be added, such as surface antigens, secreted and structural proteins and membrane receptors (other than GPCRs). The demonstrated robust performance across the protein superfamilies suggest that our attention-based approach can be extended to other superfamilies and subfamilies, provided larger enough training datasets are available for target activities.

## CONFLICTS OF INTEREST

The authors declare no conflicts of interest.

## ACKNOWLEDGMENTS

The work was funded by the Academy of Finland (No. 351507 to ZT), and Academy of Finland (grants 340141, 344698 and 345803 to TA), Norwegian Health Authority South-East (grant 2020026), the Cancer Society of Finland, and the Sigrid Jusélius Foundation. Project was also partly funded by REMEDi4ALL. The REMEDi4ALL project has received funding from the European Union’s Horizon Europe Research & Innovation program under grant agreement No 101057442. Views and opinions expressed are however those of the author(s) only and do not necessarily reflect those of the European Union or the Health and Digital Executive Agency. Neither the European Union nor the granting authority can be held responsible for them.

## Notes

### Competing Interest Statement

The authors have declared no competing interest.

## REFERENCES

[1] A. Schuhmacher, M. Hinder, A. Von Stegmann Und Stein, D. Hartl, and O. Gassmann, “Analysis of pharma R&D productivity – a new perspective needed,” Drug Discov. Today, vol. 28, no. 10, p. 103726, Oct. 2023, doi: 10.1016/j.drudis.2023.103726.

[2] R. L. Beijersbergen, “Old drugs with new tricks,” Nat. Cancer, vol. 1, no. 2, pp. 153–155, Jan. 2020, doi: 10.1038/s43018-020-0024-8.

[3] A. Gaulton et al., “ChEMBL: a large-scale bioactivity database for drug discovery,” Nucleic Acids Res., vol. 40, no. D1, pp. D1100–D1107, Jan. 2012, doi: 10.1093/nar/gkr777.

[4] T. Liu, Y. Lin, X. Wen, R. N. Jorissen, and M. K. Gilson, “BindingDB: a web-accessible database of experimentally determined protein-ligand binding affinities,” Nucleic Acids Res., vol. 35, no. Database, pp. D198–D201, Jan. 2007, doi: 10.1093/nar/gkl999.

[5] D. S. Wishart, “DrugBank: a comprehensive resource for in silico drug discovery and exploration,” Nucleic Acids Res., vol. 34, no. 90001, pp. D668–D672, Jan. 2006, doi: 10.1093/nar/gkj067.

[6] S. D. Harding et al., “The IUPHAR/BPS Guide to PHARMACOLOGY in 2024,” Nucleic Acids Res., vol. 52, no. D1, pp. D1438–D1449, Jan. 2024, doi: 10.1093/nar/gkad944.

[7] Z. Tanoli et al., “Drug Target Commons 2.0: a community platform for systematic analysis of drug– target interaction profiles,” Database, vol. 2018, Jan. 2018, doi: 10.1093/database/bay083.

[8] K. C. Cotto et al., “DGIdb 3.0: a redesign and expansion of the drug–gene interaction database,” Nucleic Acids Res., vol. 46, no. D1, pp. D1068–D1073, Jan. 2018, doi: 10.1093/nar/gkx1143.

[9] K. Han et al., “A Review of Approaches for Predicting Drug–Drug Interactions Based on Machine Learning,” Front. Pharmacol., vol. 12, p. 814858, Jan. 2022, doi: 10.3389/fphar.2021.814858.

[10] A. Vaswani, et al., “Attention Is All You Need,” 2017, doi: 10.48550/ARXIV.1706.03762.

[11] K. Huang, C. Xiao, L. M. Glass, and J. Sun, “MolTrans: Molecular Interaction Transformer for drug– target interaction prediction,” Bioinformatics, vol. 37, no. 6, pp. 830–836, May 2021, doi: 10.1093/bioinformatics/btaa880.

[12] H. Kang, S. Goo, H. Lee, J. Chae, H. Yun, and S. Jung, “Fine-tuning of BERT Model to Accurately Predict Drug–Target Interactions,” Pharmaceutics, vol. 14, no. 8, p. 1710, Aug. 2022, doi: 10.3390/pharmaceutics14081710.

[13] J. Devlin, M.-W. Chang, K. Lee, and K. Toutanova, “BERT: Pre-training of Deep Bidirectional Transformers for Language Understanding,” 2018, doi: 10.48550/ARXIV.1810.04805.

[14] Y. Kalakoti, S. Yadav, and D. Sundar, “TransDTI: Transformer-Based Language Models for Estimating DTIs and Building a Drug Recommendation Workflow,” ACS Omega, vol. 7, no. 3, pp. 2706–2717, Jan. 2022, doi: 10.1021/acsomega.1c05203.

[15] Z. Lin et al., “Evolutionary-scale prediction of atomic-level protein structure with a language model,” Science, vol. 379, no. 6637, pp. 1123–1130, Mar. 2023, doi: 10.1126/science.ade2574.

[16] J. Jumper et al., “Highly accurate protein structure prediction with AlphaFold,” Nature, vol. 596, no. 7873, pp. 583–589, Aug. 2021, doi: 10.1038/s41586-021-03819-2.

[17] N. R. C. Monteiro, J. L. Oliveira, and J. P. Arrais, “DTITR: End-to-end drug–target binding affinity prediction with transformers,” Comput. Biol. Med., vol. 147, p. 105772, Aug. 2022, doi: 10.1016/j.compbiomed.2022.105772.

[18] J. Bian, X. Zhang, X. Zhang, D. Xu, and G. Wang, “MCANet: shared-weight-based MultiheadCrossAttention network for drug–target interaction prediction,” Brief. Bioinform., vol. 24, no. 2, p. bbad082, Mar. 2023, doi: 10.1093/bib/bbad082.

[19] G. Wang et al., “Multi-TransDTI: Transformer for Drug–Target Interaction Prediction Based on Simple Universal Dictionaries with Multi-View Strategy,” Biomolecules, vol. 12, no. 5, p. 644, Apr. 2022, doi: 10.3390/biom12050644.

[20] R. Zhang, Z. Wang, X. Wang, Z. Meng, and W. Cui, “MHTAN-DTI: Metapath-based hierarchical transformer and attention network for drug–target interaction prediction,” Brief. Bioinform., vol. 24, no. 2, p. bbad079, Mar. 2023, doi: 10.1093/bib/bbad079.

[21] H. Wang, F. Guo, M. Du, G. Wang, and C. Cao, “A novel method for drug-target interaction prediction based on graph transformers model,” BMC Bioinformatics, vol. 23, no. 1, p. 459, Nov. 2022, doi: 10.1186/s12859-022-04812-w.

[22] Y. Gorishniy, I. Rubachev, and A. Babenko, “On Embeddings for Numerical Features in Tabular Deep Learning,” 2022, doi: 10.48550/ARXIV.2203.05556.

[23] Z. Tanoli, Z. Alam, A. Ianevski, K. Wennerberg, M. Vähä-Koskela, and T. Aittokallio, “Interactive visual analysis of drug–target interaction networks using Drug Target Profiler, with applications to precision medicine and drug repurposing,” Brief. Bioinform., Dec. 2018, doi: 10.1093/bib/bby119.

[24] E. F. Douglass et al., “A community challenge for a pancancer drug mechanism of action inference from perturbational profile data,” Cell Rep. Med., vol. 3, no. 1, p. 100492, Jan. 2022, doi: 10.1016/j.xcrm.2021.100492.

[25] Y. Gorishniy, I. Rubachev, V. Khrulkov, and A. Babenko, “Revisiting Deep Learning Models for Tabular Data.” arXiv, Oct. 26, 2023. [Online]. Available: http://arxiv.org/abs/2106.11959

[26] A. Cereto-Massagué, M. J. Ojeda, C. Valls, M. Mulero, S. Garcia-Vallvé, and G. Pujadas, “Molecular fingerprint similarity search in virtual screening,” Methods, vol. 71, pp. 58–63, Jan. 2015, doi: 10.1016/j.ymeth.2014.08.005.

[27] A. Cichonska et al., “Learning with multiple pairwise kernels for drug bioactivity prediction,” Bioinformatics, vol. 34, no. 13, pp. i509–i518, Jul. 2018, doi: 10.1093/bioinformatics/bty277.

[28] V. Periwal et al., “Bioactivity assessment of natural compounds using machine learning models trained on target similarity between drugs,” PLOS Comput. Biol., vol. 18, no. 4, p. e1010029, Apr. 2022, doi: 10.1371/journal.pcbi.1010029.

[29] J. L. Durant, B. A. Leland, D. R. Henry, and J. G. Nourse, “Reoptimization of MDL Keys for Use in Drug Discovery,” J. Chem. Inf. Comput. Sci., vol. 42, no. 6, pp. 1273–1280, Nov. 2002, doi: 10.1021/ci010132r.

[30] G. Landrum, “RDKit Documentation.” May 13, 2019. [Online]. Available: https://buildmedia.readthedocs.org/media/pdf/rdkit/latest/rdkit.pdf

[31] P. Gedeck, B. Rohde, and C. Bartels, “QSAR − How Good Is It in Practice? Comparison of Descriptor Sets on an Unbiased Cross Section of Corporate Data Sets,” J. Chem. Inf. Model., vol. 46, no. 5, pp. 1924–1936, Sep. 2006, doi: 10.1021/ci050413p.

[32] D. Rogers and M. Hahn, “Extended-Connectivity Fingerprints,” J. Chem. Inf. Model., vol. 50, no. 5, pp. 742–754, May 2010, doi: 10.1021/ci100050t.

[33] Z.-X. Yue et al., “A systematic review on the state-of-the-art strategies for protein representation,” Comput. Biol. Med., vol. 152, p. 106440, Jan. 2023, doi: 10.1016/j.compbiomed.2022.106440.

[34] F. Langenfeld et al., “SHREC 2020: Multi-domain protein shape retrieval challenge,” Comput. Graph., vol. 91, pp. 189–198, Oct. 2020, doi: 10.1016/j.cag.2020.07.013.

[35] M. Novotni and R. Klein, “Shape retrieval using 3D Zernike descriptors,” Comput.-Aided Des., vol. 36, no. 11, pp. 1047–1062, Sep. 2004, doi: 10.1016/j.cad.2004.01.005.

[36] A. Cichonska et al., “Computational-experimental approach to drug-target interaction mapping: A case study on kinase inhibitors,” PLOS Comput. Biol., vol. 13, no. 8, p. e1005678, Aug. 2017, doi: 10.1371/journal.pcbi.1005678.

[37] D. Kihara, L. Sael, R. Chikhi, and J. Esquivel-Rodriguez, “Molecular Surface Representation Using 3D Zernike Descriptors for Protein Shape Comparison and Docking,” Curr. Protein Pept. Sci., vol. 12, no. 6, pp. 520–530, Sep. 2011, doi: 10.2174/138920311796957612.

[38] L. Di Rienzo, L. De Flaviis, G. Ruocco, V. Folli, and E. Milanetti, “Binding site identification of G protein-coupled receptors through a 3D Zernike polynomials-based method: application to C. elegans olfactory receptors,” J. Comput. Aided Mol. Des., vol. 36, no. 1, pp. 11–24, Jan. 2022, doi: 10.1007/s10822-021-00434-1.

[39] Y. Saeys, I. Inza, and P. Larrañaga, “A review of feature selection techniques in bioinformatics,” Bioinformatics, vol. 23, no. 19, pp. 2507–2517, Oct. 2007, doi: 10.1093/bioinformatics/btm344.

[40] A. I. Cowen-Rivers et al., “HEBO: An Empirical Study of Assumptions in Bayesian Optimisation,” J. Artif. Intell. Res., vol. 74, pp. 1269–1349, Jul. 2022, doi: 10.1613/jair.1.13643.

[41] R. Turner et al., “Bayesian Optimization is Superior to Random Search for Machine Learning Hyperparameter Tuning: Analysis of the Black-Box Optimization Challenge 2020.” arXiv, Aug. 31, 2021. [Online]. Available: http://arxiv.org/abs/2104.10201

[42] L. Wenlong, Y. Fan, Y. Changhao, Z. Dian, and Z. Xuan, “Batch Bayesian Optimization via Multi-objective Acquisition Ensemble for Automated Analog Circuit Design,” vol. 80, pp. 3306–3314, 2018.

[43] L. Li, K. Jamieson, G. DeSalvo, A. Rostamizadeh, and A. Talwalkar, “Hyperband: A Novel Bandit-Based Approach to Hyperparameter Optimization,” 2016, doi: 10.48550/ARXIV.1603.06560.

[44] J. Tan, J. Yang, S. Wu, G. Chen, and J. Zhao, “A critical look at the current train/test split in machine learning.” arXiv, Jun. 08, 2021. [Online]. Available: http://arxiv.org/abs/2106.04525

[45] M. I. Davis et al., “Comprehensive analysis of kinase inhibitor selectivity,” Nat. Biotechnol., vol. 29, no. 11, pp. 1046–1051, Nov. 2011, doi: 10.1038/nbt.1990.

[46] T. Pahikkala et al., “Toward more realistic drug-target interaction predictions,” Brief. Bioinform., vol. 16, no. 2, pp. 325–337, Mar. 2015, doi: 10.1093/bib/bbu010.

[47] T. Nguyen, H. Le, T. P. Quinn, T. Nguyen, T. D. Le, and S. Venkatesh, “GraphDTA: predicting drug–target binding affinity with graph neural networks,” Bioinformatics, vol. 37, no. 8, pp. 1140– 1147, May 2021, doi: 10.1093/bioinformatics/btaa921.

[48] T. He, M. Heidemeyer, F. Ban, A. Cherkasov, and M. Ester, “SimBoost: a read-across approach for predicting drug–target binding affinities using gradient boosting machines,” J. Cheminformatics, vol. 9, no. 1, p. 24, Dec. 2017, doi: 10.1186/s13321-017-0209-z.

[49] J. Shim, Z.-Y. Hong, I. Sohn, and C. Hwang, “Prediction of drug–target binding affinity using similarity-based convolutional neural network,” Sci. Rep., vol. 11, no. 1, p. 4416, Feb. 2021, doi: 10.1038/s41598-021-83679-y.

[50] H. Öztürk, A. Özgür, and E. Ozkirimli, “DeepDTA: deep drug–target binding affinity prediction,” Bioinformatics, vol. 34, no. 17, pp. i821–i829, Sep. 2018, doi: 10.1093/bioinformatics/bty593.

[51] K. Abbasi, P. Razzaghi, A. Poso, M. Amanlou, J. B. Ghasemi, and A. Masoudi-Nejad, “DeepCDA: deep cross-domain compound–protein affinity prediction through LSTM and convolutional neural networks,” Bioinformatics, vol. 36, no. 17, pp. 4633–4642, Nov. 2020, doi: 10.1093/bioinformatics/btaa544.

