## Supplementary figures and images for "Attention-based approach to predict drug-target interactions across seven target superfamilies"

### Supplemental Data 1

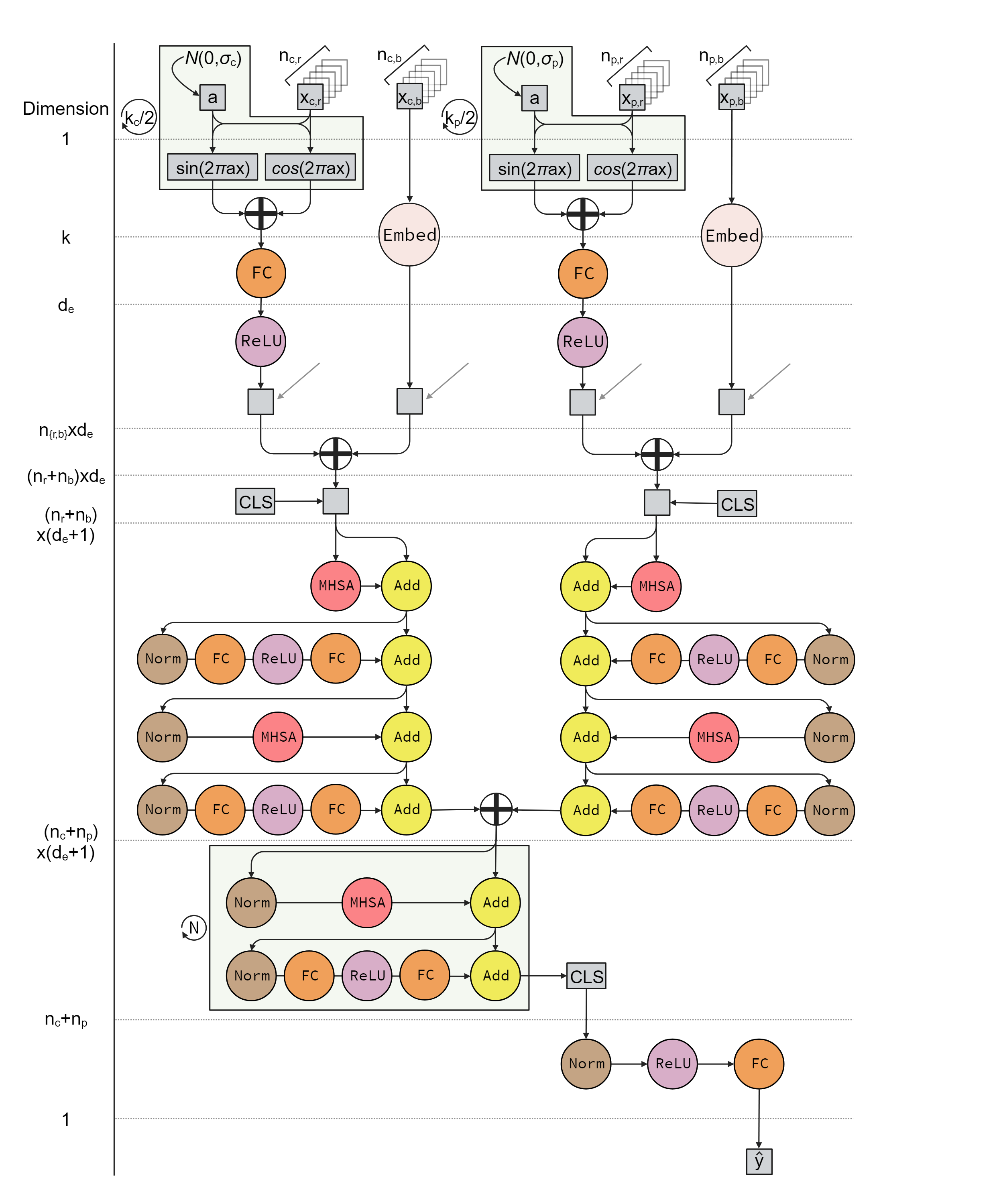

### Supplemental Data 2

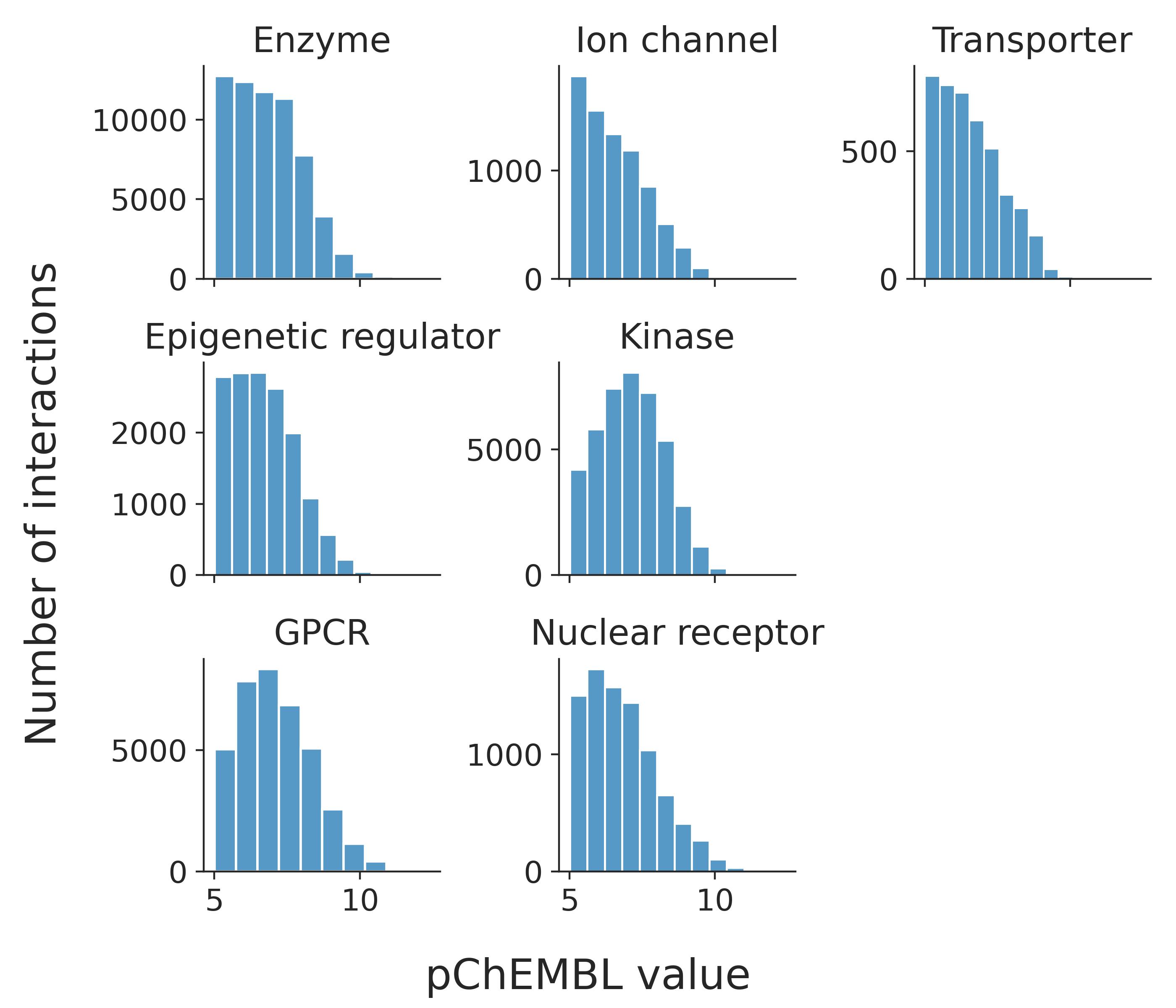

### Supplemental Data 3

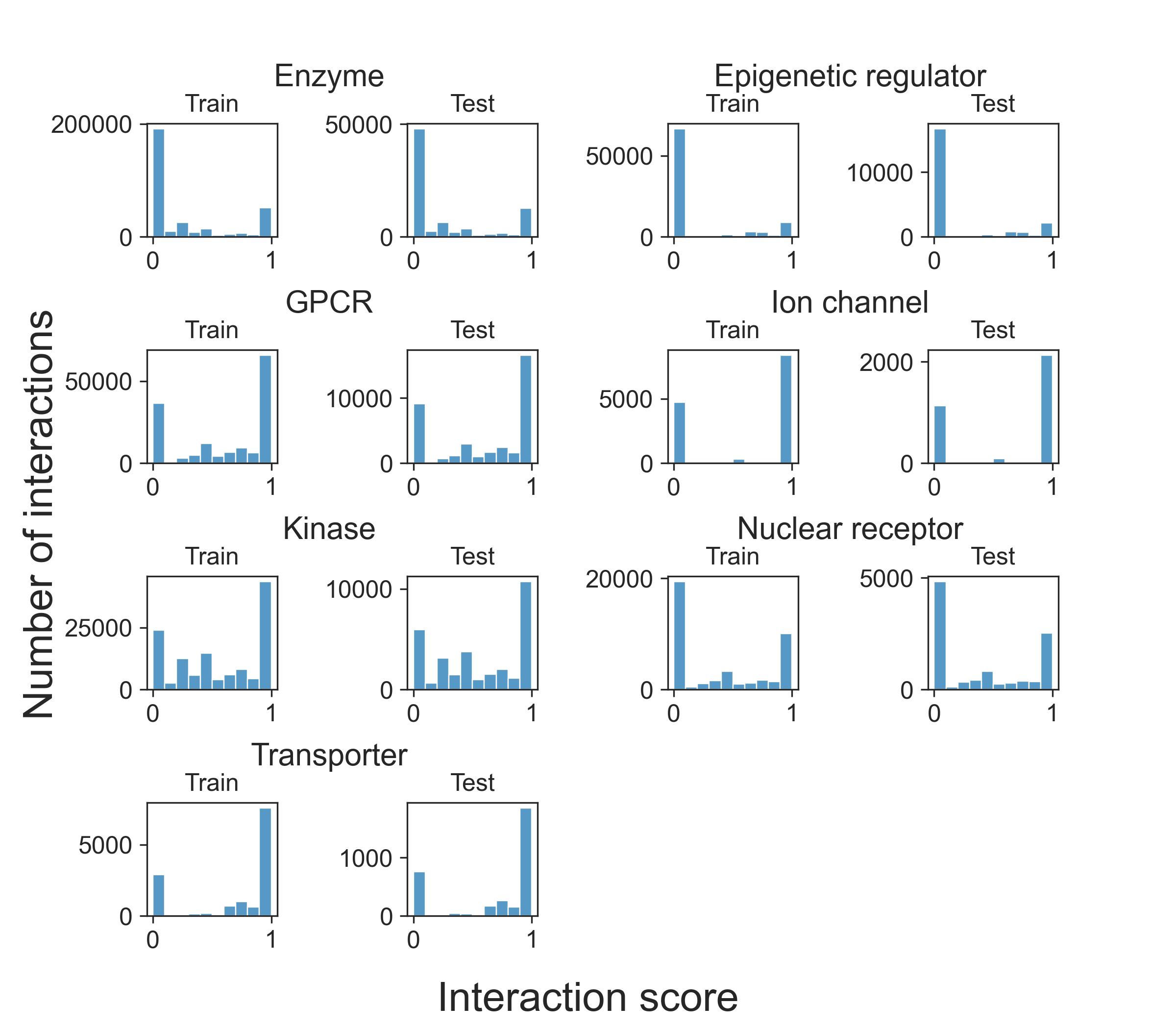

### Supplemental Data 4

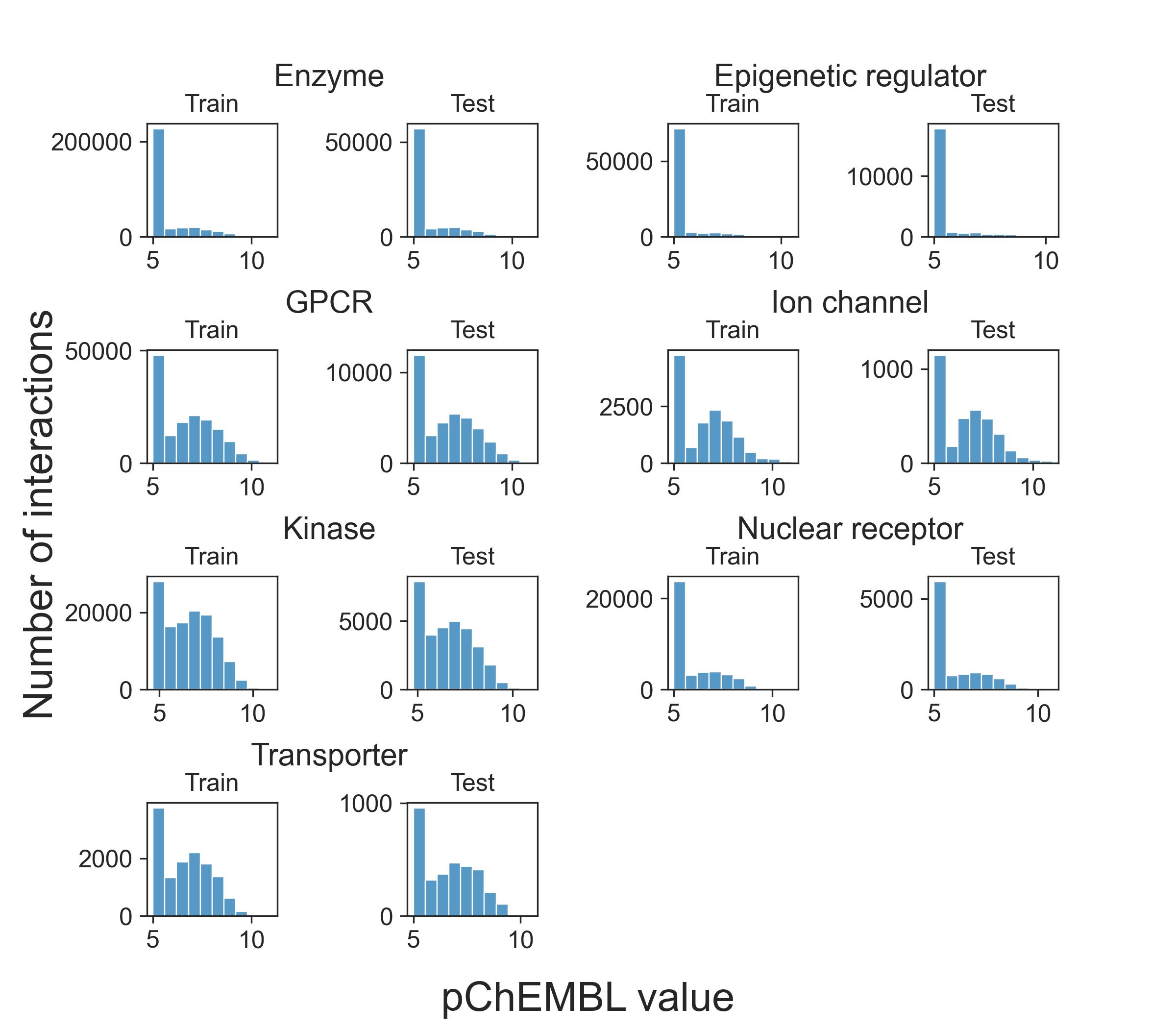
